# Real-Time Computation of Brain E-Field for Enhanced Transcranial Magnetic Stimulation Neuronavigation and Optimization

**DOI:** 10.1101/2023.10.25.564044

**Authors:** Nahian I. Hasan, Moritz Dannhauer, Dezhi Wang, Zhi-De Deng, Luis J. Gomez

**Author notes:** (N.I. Hasan); (M. Dannhauer); (D. Wang); (Z. Deng); (L.J. Gomez).

## Abstract

Transcranial Magnetic Stimulation (TMS) coil placement and pulse waveform current are often chosen to achieve a specified E-field dose on targeted brain regions. TMS neuronavigation could be improved by including real-time accurate distributions of the E-field dose on the cortex. We introduce a method and develop software for computing brain E-field distributions in real-time enabling easy integration into neuronavigation and with the same accuracy as 1^st^-order finite element method (FEM) solvers. Initially, a spanning basis set (< 400) of E-fields generated by white noise magnetic currents on a surface separating the head and permissible coil placements are orthogonalized to generate the modes. Subsequently, Reciprocity and Huygens’ principles are utilized to compute fields induced by the modes on a surface separating the head and coil by FEM, which are used in conjunction with online (real-time) computed primary fields on the separating surface to evaluate the mode expansion. We conducted a comparative analysis of E-fields computed by FEM and in real-time for eight subjects, utilizing two head model types (SimNIBS’s ‘headreco’ and ‘mri2mesh’ pipeline), three coil types (circular, double-cone, and Figure-8), and 1000 coil placements (48,000 simulations). The real-time computation for any coil placement is within 4 milliseconds (ms), for 400 modes, and requires less than 4 GB of memory on a GPU. Our solver is capable of computing E-fields within 4 ms, making it a practical approach for integrating E-field information into the neuronavigation systems without imposing a significant overhead on frame generation (20 and 50 frames per second within 50 and 20 ms, respectively).

## 1. Introduction

Transcranial magnetic stimulation (TMS) is a non-invasive brain stimulation approach that is widely used in neuroscience research to study brain function and has been approved by Food and Drug Administration (FDA) in the treatment of depression, obsessive-compulsive disorder, migraine, and smoking cessation [1, 2, 3, 4, 5, 6, 7, 8, 9, 10, 11]. TMS uses coils driven by low-frequency current pulses to stimulate targeted brain regions. Computational electric field (E-field) dosimetry can be used to 1) quantify the E-field strength and distribution to determine brain regions affected by TMS, 2) identify coil placements and orientation that would maximize the E-field induced at a prespecified target, and 3) design coils with a desired E-field profile [12, 13, 14, 15, 16, 17, 18]. All of these applications require repeated execution of E-field solvers to determine the E-field. Correspondingly, there is ongoing interest in fast and accurate E-field solvers for TMS [12, 19, 20]. In particular, there is an interest in incorporating accurate and precise E-field information in neuronavigation systems that use subject MRI along with cameras to provide practitioners with precise information about the coil location relative to the head. Appending E-field information to neuronavigation tools would allow for on-the-fly E-field informed coil placement reconfiguration and intensity dosing to target multiple or even changing cortical targets during a single TMS session [21, 22]. The integration of E-field information into neuronavigation requires a solver that can accurately determine the E-field in real-time.

Several approaches have been proposed for determining the E-field in real time. For example, the most widely used approach locally fits a sphere to the head to estimate the E-field under the coil rapidly [23, 24]. However, this solver does not account for local neural geometry, consequently yielding less accurate local E-field estimates. Furthermore, deep learning approaches have been introduced, which have been promising. However, at this point, they still result in a high relative residual error of about 6% even in a small target region [25] and 18% in the whole brain in [26]. More recently, boundary element method (BEM)-based near real-time solvers that can account for local brain anatomy have been developed [19, 20].

Stenroos et al. (2019) used efficient quadratures for the coil and tissue boundary sources enabling real-time E-field solutions using GPUs [19]. Specifically, they use pre-computed boundary potentials on a mesh to compute the TMS-induced E-field in a cortical region of interest (ROI) in real time via reciprocity. Using their method, the computation scales as the “number of coil quadrature nodes” times “the number of cortical E-field evaluation points” times “the number of vertices of the surface meshes”.

Correspondingly, to maintain fast computation, significant trade-offs have to be made between mesh resolution and the number of ROI points. For example, to achieve 36 ms on a GPU they used a mesh consisting of 21,052 nodes and 20,324 cortical ROI locations. The Ernie mesh, a typical head mesh generated by SimNIBS software v3.2, has more than 30 times the number of mesh nodes (667,000 nodes) and more than 10 times the number of cortical surface nodes (216,130 nodes); this would result in a 300 fold computation time. Furthermore, newer more detailed head models generated using SimNIBS v4.0 contain an even higher number of nodes. As a result, to achieve real-time this approach is limited to much lower fidelity head models than what is typically used.

To overcome these challenges, Daneshzand et al. proposed the Magnetic Stimulation Profile (MSP) approach that approximates the TMS coil-induced E-fields in a cortical ROI as a linear combination of dipole-induced E-fields [20]. Dipole-induced E-fields are precomputed in a process that can take from 5 to 18 hours depending on desired accuracy and mesh resolution. Then, these precomputed E-fields are used in real-time to determine the TMS coil-induced E-fields. To find E-field expansion coefficients the method employs a least squares to match the primary E-field of the coil and the primary E-fields induced by a linear combination of dipoles. This method requires ∼3000 dipole E-fields, requiring ∼ 32 GBs of memory and about 0.37 s using a Xeon E5-2360 CPU for 120,000 cortical triangles to match the full BEM solution with an average relative residual error of about 10% and a 5% error in the predicting the peak E-field [20]. The memory and CPU time of this method scales as the number of dipole E-fields times the number of E-field evaluation points. As such, to lower the memory requirements and achieve real-time performance, significant trade-offs between accuracy and cortical E-field samples have to be made.

In this paper, similar to the MSP approach, we approximate the E-field as a linear combination of precomputed E-fields. However, to address the computational bottlenecks, we use a novel method for determining an E-field basis in lieu of the dipole E-field basis. This results in an accurate and robust real-time E-field solver for any type of TMS coils. To find an efficient basis of E-field modes in the cortex due to any source outside of the head we apply an approach similar to the probabilistic matrix decomposition (PMD) [27] based approach. By using modes in lieu of dipole E-fields, we reduce the required number of modes more than tenfold relative to the MSP approach. For example, to reach a 10% error we require 110 modes relative to the 3,000 modes required using the MSP approach. Furthermore, our approach requires under 400 basis modes to estimate the TMS-induced E-field with an error lower than 3%. To rapidly determine the expansion coefficients, we introduce reciprocity and Huygens’ principle formalism that enables us to relate the primary E-fields and magnetic field (H-field) on a fictitious surface engulfing the head to expansion coefficients. Using this method, we are able to estimate the E-field to 2% error within 4 ms from the primary E-field and H-field for 216,000 cortical surface targets. Furthermore, unlike the MSP, the expansion coefficients are found in an analytical way without the need for regularization to prevent over-fitting, making this a more robust approach.

## 2. Methods

### 2.1. Overview

This section describes the procedure for real-time determination of an approximate expansion for the E-field induced by the coil in the brain during TMS

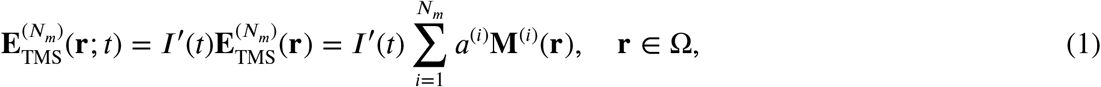

where **r** denotes a Cartesian location, Ω denotes the brain region, **M**^(*i*)^(**r**) and *a*^(*i*)^ are one of the *Nm* mode functions and expansion coefficients, respectively, *I*^′^(*t*) is time-derivative of the driving current pulse waveform, and 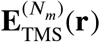 is the approximate E-field expansion at an instant when *I*^′^(*t*) is 1. Note that in the rest of this paper, we adopt quasi-stationary assumptions that have been shown valid and are standard for TMS E-field modeling [12, 13, 20, 28, 29, 30]. Correspondingly, it is assumed that the H-fields and E-fields are proportional to the coil driving current and its temporal derivative, respectively. As such, their temporal dependence is suppressed in all equations that follow.

In Section 2.2, we describe a procedure for finding orthonormal mode functions **M**^(*i*)^(**r**) (i.e., *⟨***M**^(*i*)^, **M**^(*j*)^⟩ = *δ*_*i,j*_, where *⟨***f**, **g**⟩ = ∫Ω **f** (**r**) ⋅ **g**(**r**)d**r** and *δ*_*i,j*_ is the Kronecker delta function) that can efficiently represent the E-fields induced in the brain by any TMS coil.

Once these functions are determined, the coefficients *a*^(*i*)^ are chosen to minimize the *L*^2^ error of the expansion (i.e., argmin 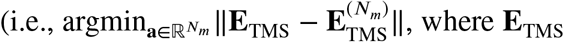, where **E**is the actual E-field induced during TMS,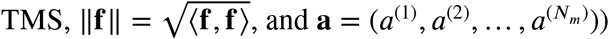, and **a** = (*a*^(1)^, *a*^(2)^, …, *a*^(*N*_*m*_^))).

Since the mode functions are orthonormal, the coefficients are *a*^(*i*)^ = *⟨***M**^(*i*)^, **E**_TMS_ ⟩. The above expression for determining each *a*^(*i*)^ requires *a priori* knowledge of the TMS-induced E-field in the brain. As such, it is not amenable to real-time solvers. Alternatively, in Section 2.3, we use reciprocity and Huygens’ equivalence principles to show that

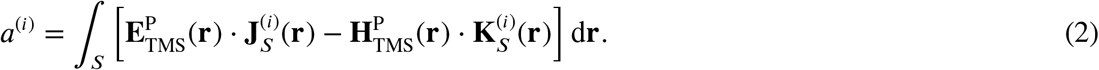

Here, *S* denotes the Huygens’ surface separating the head and the coil and is to be defined in Section 2.3, 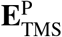 and 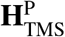 are the primary E-field and H-field due to the coil evaluated on Huygens’ surface and 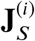 and 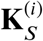 are equivalent surface currents associated with the *i*^th^ mode function. Eq. (2) only requires primary fields on Huygens’ surface and here it is used to determine expansion coefficients in real-time. In Section 2.3, we provide a method for determining 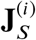 and 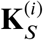 from **M**^(*i*)^ and evaluating (2) via quadrature. Finally, in Section 2.4, we provide a method for rapidly determining 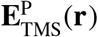 (**r**) and 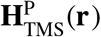 (**r**) on the Huygens’ surface.

### 2.2. Generation of mode function M^(*i*)^

The mode functions **M**^(*i*)^(**r**), where *i* = 1, 2, …, *N*_*m*_, are derived by finding an orthonormal basis to the E-fields induced in the brain by *N*_*m*_ magnetic surface currents residing on a fictitious surface 1 mm directly outside of the scalp (Fig. 1). The *N*_*m*_ current distributions are chosen as distinct realizations of Gaussian white noise. The *N*_*m*_ Gaussian white noise magnetic current realizations are continuous analogous to the Gaussian white noise vectors that we previously successfully used to compress matrices of TMS-induced brain E-field samples for many coil placements and brain locations [31].

**Figure 1:**
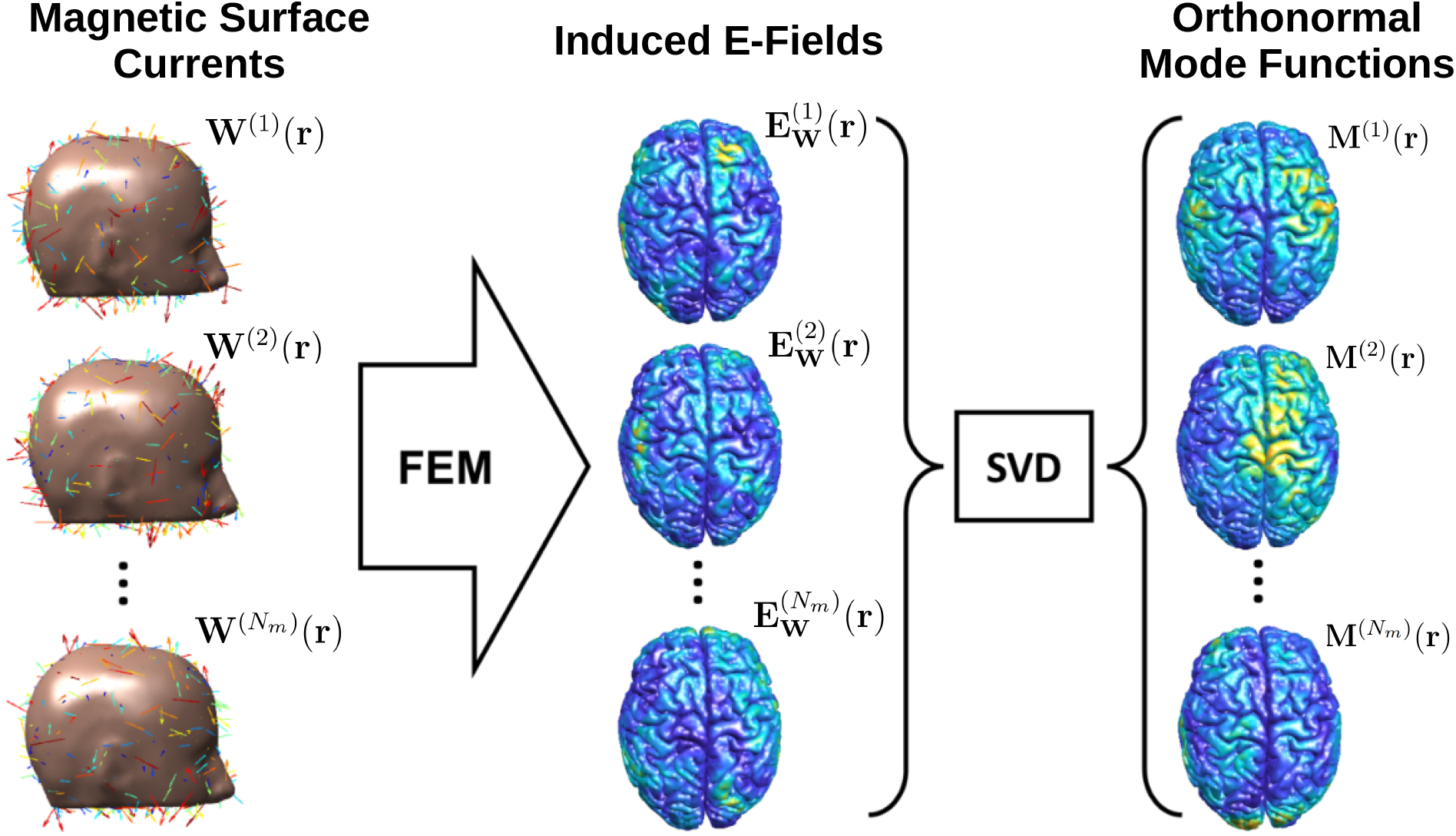
Generation of Mode Functions from surface magnetic currents. The left column shows the individual realization of magnetic surface currents (**W**^(*i*)^(**r**)). The middle column shows the induced E-field on the brain for each surface current distribution, generated by an FEM simulation. The right column shows the *N*_*m*_ orthonormal mode functions [**M**^(*i*)^(**r**), *i* = 1, 2, …, *N*_*m*_], generated by a singular value decomposition (SVD) over the *N*_*m*_ induced E-fields.

The fictitious surface is first approximated by a triangle mesh consisting of *Nd* triangles to approximate the E-fields generated by these currents. The triangle mesh is generated by extruding the nodes of the scalp surface mesh 1 mm normally outward. The E-fields are then determined by equivalent magnetic dipole moments generated by sampling the magnetic surface currents at the center of each triangle and multiplying them with the area of the respective triangle. With the above approximation, the E-field generated by the *i*^th^ realization of the white noise distributed magnetic surface current in free space (i.e., the primary E-field) is

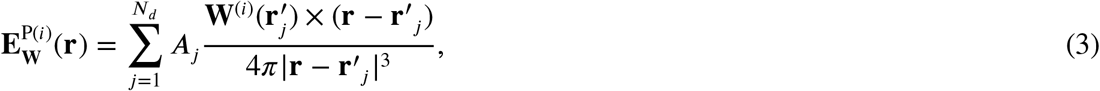

where *A*_*j*_ and 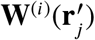are the area and the *i*^th^ magnetic surface current at the center of the *j*^th^ triangle, respectively. The value of 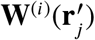 is determined using a normally distributed random number generator.

The total E-field induced in the brain by current **W**^(*i*)^(**r**) is 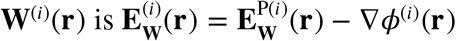, where *ϕ*^(*i*)^(**r**) is the scalar potential. The scalar potential is determined by solving the current continuity equation

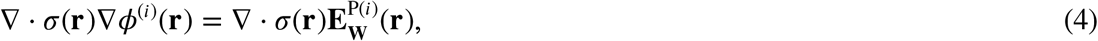

where σ(**r**) is the conductivity at location **r** in the head. The region surrounding the head is assumed to be insulating (i.e., σ(**r**) = 0). Correspondingly, 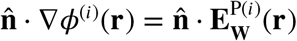 on the scalp boundary. Here ñ is a normal vector on the scalp surface pointing outward. To solve Eq. (4), we approximate the head by a tetrahedral mesh where each tetrahedron is assigned a homogeneous conductivity.

Then, Eq. (4) is solved using either our first or second-order in-house finite element solvers that have been validated [30] and available online [32].

The total E-field in the brain is approximated as a piece-wise constant within each tetrahedron.

The resulting expansion is 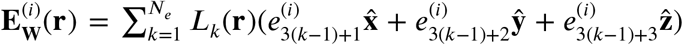, where *N*e is the number of tetrahedrons in the brain and *Lk*(**r**) = 1 in the *k*^th^ tetrahedron and zero outside it.

To find the set of orthonormal mode functions **M**^(*i*)^(**r**), where *i* = 1, 2, …, *N*_*m*_, from the E-field results we must orthogonalize them with respect to their inner-product

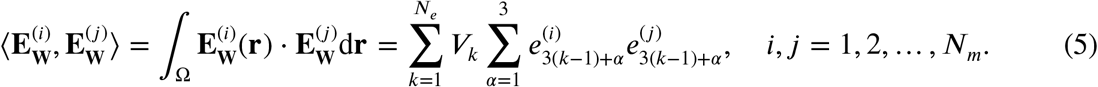

The above establishes an equivalence between the inner-product of two E-fields and the dot-product of two column vectors of a matrix **Z** with entries 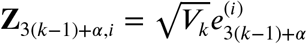, where *k* = 1, 2, …, *N*_e_, α = 1, 2, 3, *i* = 1, 2, …, *N*_*m*_, and **V**_*k*_ is the volume of the *k*^th^ tetrahedron. To find orthonormal mode functions from **Z** we first compute its singular value decomposition (SVD) **Z** = **USV**^T^. The modes are defined as

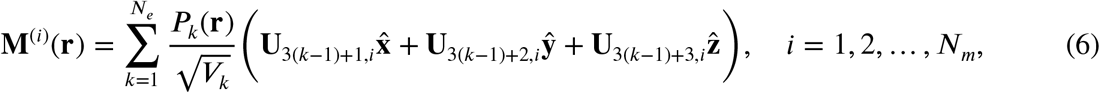

where **U**_3(*k*−1)+α,*i*_ is the *i*^th^ column and (3(*k* − 1) + α)^th^ row entry of **U**.

### 2.3. Evaluation of coefficients *a*^(*i*)^

In this section, we provide proof of Eq. (2) and details of the numerical implementation used to compute the coefficients *a*^(*i*)^.

We first apply the reciprocity principle to relate two scenarios: one where the TMS coil induces E-fields in the head and another where cortical impressed currents in the head (**M**^(*i*)^(**r**)) induces E-fields outside the head [Fig. 2 (A) and (B)]. Specifically, the reciprocity principle establishes the following

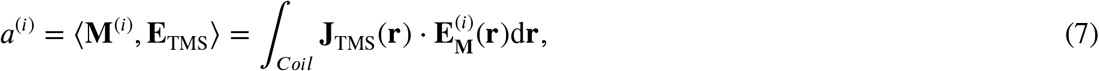

where 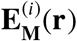 is the E-field generated by impressed currents **M**^(*i*)^(**r**), and *Coil* is the coil support.

**Figure 2:**
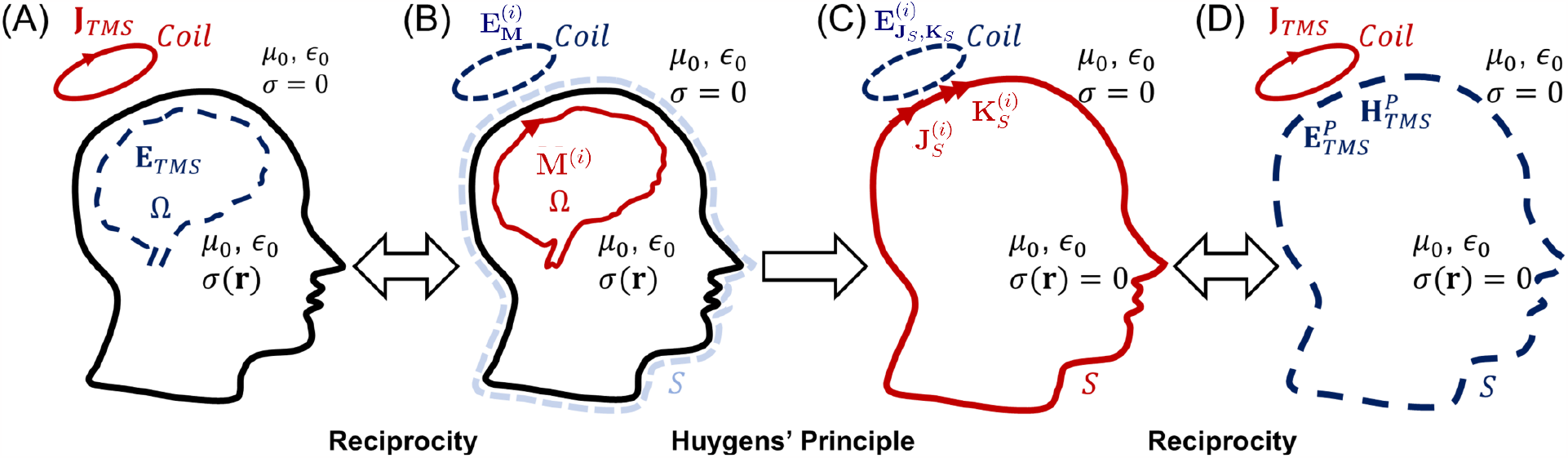
(A) The expansion coefficients can be computed from the induced E-field in the brain due to coil current outside the scalp. (B) Electromagnetic Reciprocity dictates that the expansion coefficients can also be computed by determining the E-field induced on the coil by mode sources in the brain. (C) According to Huygens’ principle, the fields outside the head generated by the mode sources in the brain can be represented as arising from equivalent electric and magnetic currents on the Huygens’ surface radiating in space. (D) Reciprocity dictates that the expansion coefficients can be computed from the primary E-fields and H-fields on the Huygens’ surface induced by the coil.

**Figure 3:**
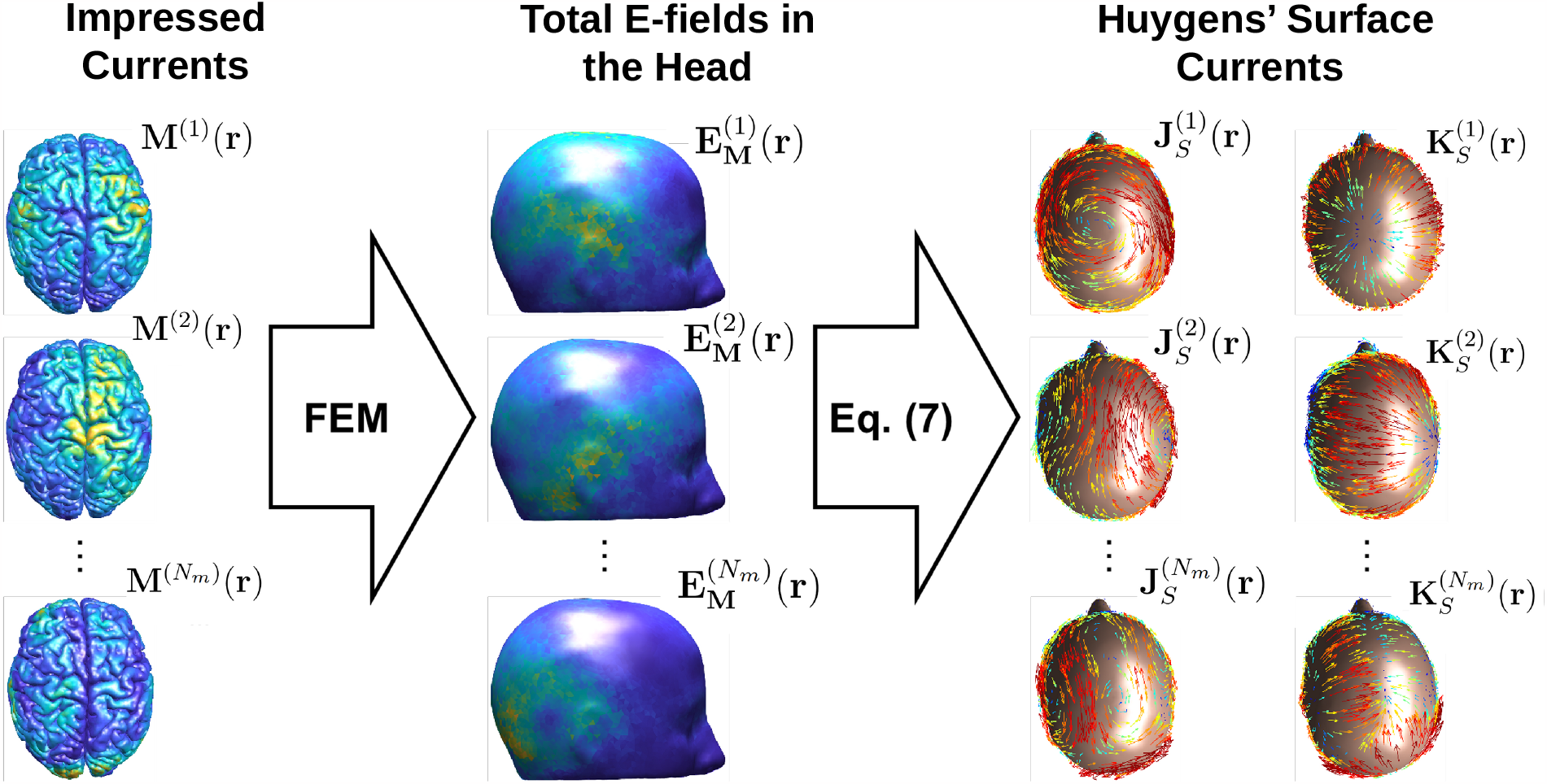
E-fields generated by the individual realization of the orthonormal mode functions or impressed currents (**M**^(*i*)^(**r**)) are evaluated on the Huygens’ surface. Then the electric and magnetic currents are calculated using the reciprocity principle on the Huygens’ surface.

Eq. (7) enables the computation of *a*^(*i*)^ from the E-fields generated outside of the head by the impressed current **M**^(*i*)^(**r**) in the brain.

By leveraging Huygens’ principle, the fields outside the head due to impressed current **M**^(*i*)^(**r**) can be represented as being generated from equivalent surface electric currents 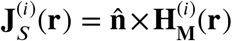 and magnetic currents 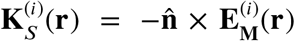 residing on a surface separating the head and coil (referred to as the Huygens’ surface or *S*) [Fig. 2 (B) and (C)]. Here **H**^(*i*)^(**r**) is the H-field generated by current **M**^(*i*)^(**r**). Both 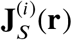 and 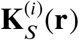 generate E-field 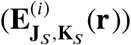 inside and outside of the Huygens’ surface. According to the Huygens’ principle, 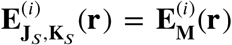. Furthermore, these equivalent currents radiate in free-space. Applying the reciprocity principle to the equivalent scenario results in Eq. (2) [Fig. 2 (C) and (D)].

To evaluate *a*^(*i*)^ numerically, we need to first determine the equivalent electric and magnetic currents on the Huygens’ surface for each of the impressed currents **M**^(*i*)^(**r**). The equivalent electric and magnetic currents on the Huygens’ surface are

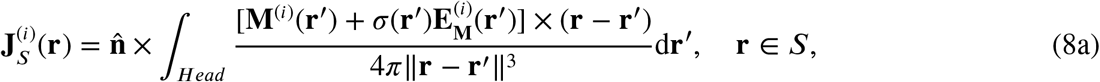

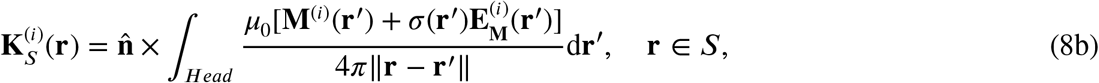

where *H*e*ad* is the whole head region.

To evaluate Eq. (8), we first need to determine 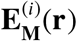 in the head (procedure shown in Fig. (3). This is done by applying a FEM procedure to ∇⋅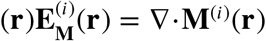 as described in [12, 13, 31] (available online at [32]). The integrals in Eq. (8) are approximated by applying a single point Gaussian quadrature rule for each head mesh tetrahedron and are rapidly evaluated using the FMM library [33].

The Huygens’ surface is chosen to be the same as the fictitious surface in Section 2.2. To determine *a*^(*i*)^, a single point Gaussian quadrature is used. This results in the following

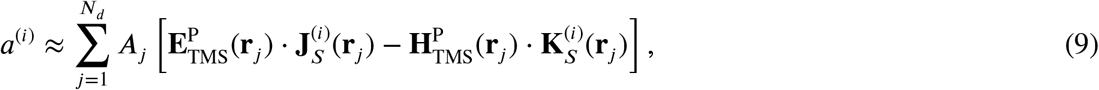

where *A*_*j*_ and **r**_*j*_ are the area and the center position of the *j*^th^ triangle.

### 2.4. Fast evaluation of primary fields due to TMS coils

Evaluation of *a*^(*i*)^ and the TMS-induced E-field requires computation of the primary fields generated by the TMS coil on the Huygens’ surface. As such, each time the coil moves these must be computed in real-time. Here, we adopt the approach proposed in [20, 29] that leverages the fact that the primary fields are functions of position relative to the coil (i.e., translational invariance). As a result, the primary fields are rapidly computed by interpolating samples on a 3D Cartesian grid using a process described next.

Here we assume a reference coil placement flat on and centered about the x-y plane. The primary fields are sampled on a 3D grid with 4 mm grid spacing with 822,000 grid points (Fig. 4). This grid spacing empirically was found to result in an interpolation error of the order of 10^−3^%. Results of interpolation errors for various grid spacing are given in the supplementary (Fig. S1).

**Figure 4:**
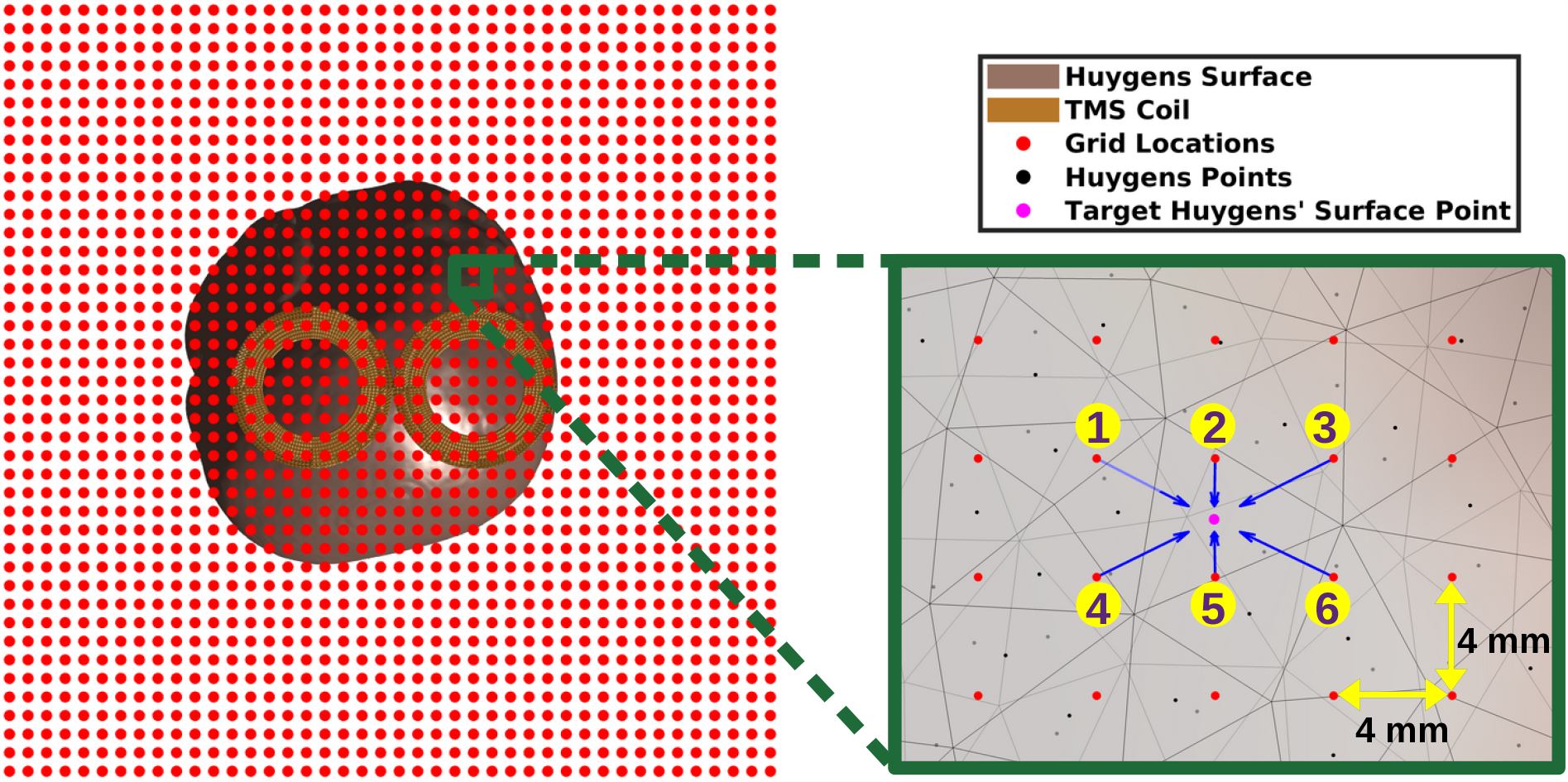
Inverse Relative transformation of the Huygens surface with respect to the TMS coil inside the interpolation grid points (red). The right figure shows an illustration of the multi-linear interpolation process for an exemplary targeted Huygens’ surface node (pink), where the primary field is interpolated by the nearby grid points (numbered 1–6).

The grid is large enough to contain the whole Huygens’ surface for any relative placement with respect to the coil. Coil placements are defined as coordinate transformation (i.e., a rotation and translation) to the coil relative to the reference coil placement as

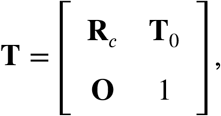

where **R**_*c*_ is a 3 × 3 rotation matrix and **T**_0_ is a 3 × 1 translation vector. Applying the transformation **T** to the coil is equivalent to applying **T**^−1^ to the Huygens’ surface. We apply the transformation **T**^−1^ to the subjects’ Huygens’ surface instead of the transformation **T** to the coil. This avoids the need to translate the grid that has 822, 000 grid points and only requires translating 120, 000 Huygens’ surface nodes. Furthermore, this enables us to use the computationally efficient standard multi-linear interpolation to generate E-field samples at the centers of the Huygens’ surface triangular facets.

### 2.5. Summary of the real-time TMS pipeline

In this section, we summarize the offline mode and surface equivalent current calculation stage and the real-time E-field calculation stage. Algorithm 1 summarizes the critical steps for computing the mode functions. Algorithm 2 describes the four fundamental steps to calculate the TMS-induced E-field in the ROI in real time while the modes and primary fields are already pre-computed.

### 2.6. Coil and head models

The algorithm is tested on fourteen MRI-derived heads and three distinct TMS coil models. The MRI-derived head models are generated from 8 distinct subject MRIs. One is the ‘Ernie’ subject included in SimNIBS-3.2 [34] and the 7 others were collected from [35]. The fourteen head models are generated using the ‘mri2mesh’ and ‘headreco’ tools in SimNIBS [36, 37]. The ‘mri2mesh’ models comprise 668,000–742,000 nodes and 3.73–4.16 million tetrahedrons. On the other hand, the ‘headreco’ models consist of 528,000–886,000 nodes and 2.87–4.92 million tetrahedrons.

We consider only the five homogeneous concentric compartments such as (from inner to outer) white matter (WM), grey matter (GM), cerebrospinal fluid (CSF), skull, and scalp with corresponding conductivity of 0.126, 0.275, 1.654, 0.01, and 0.465 S⁄m, respectively [38] in all head models. The time required for the generation of each head model was between 20–24 hours using the ‘mri2mesh’ tool and 1.5–2 hours using the ‘headreco’ tool.

#### Algorithm 1 Pre-processing stage (Mode and equivalent surface current calculation)

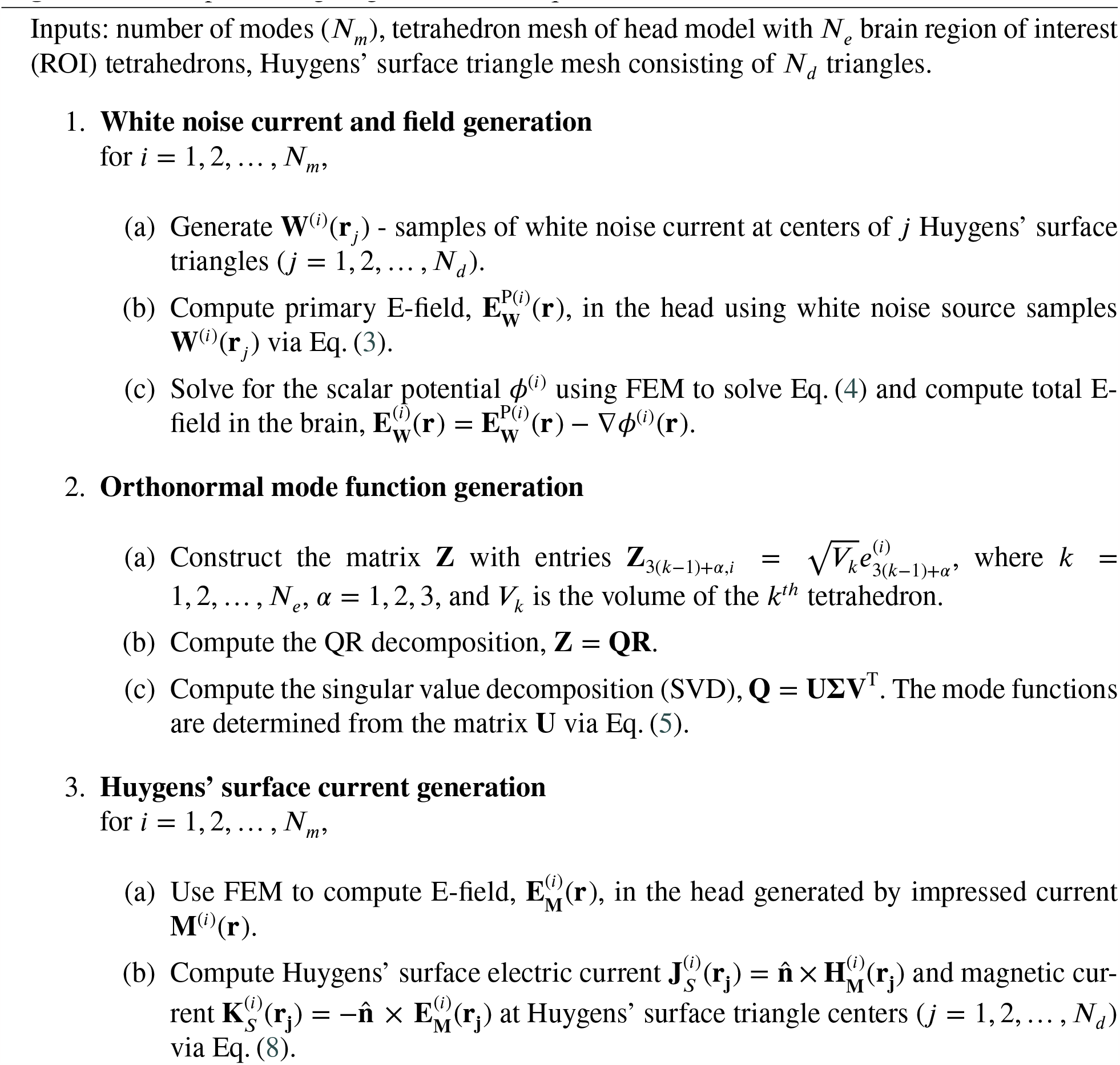

#### Algorithm 2 Real-Time E-field Calculation

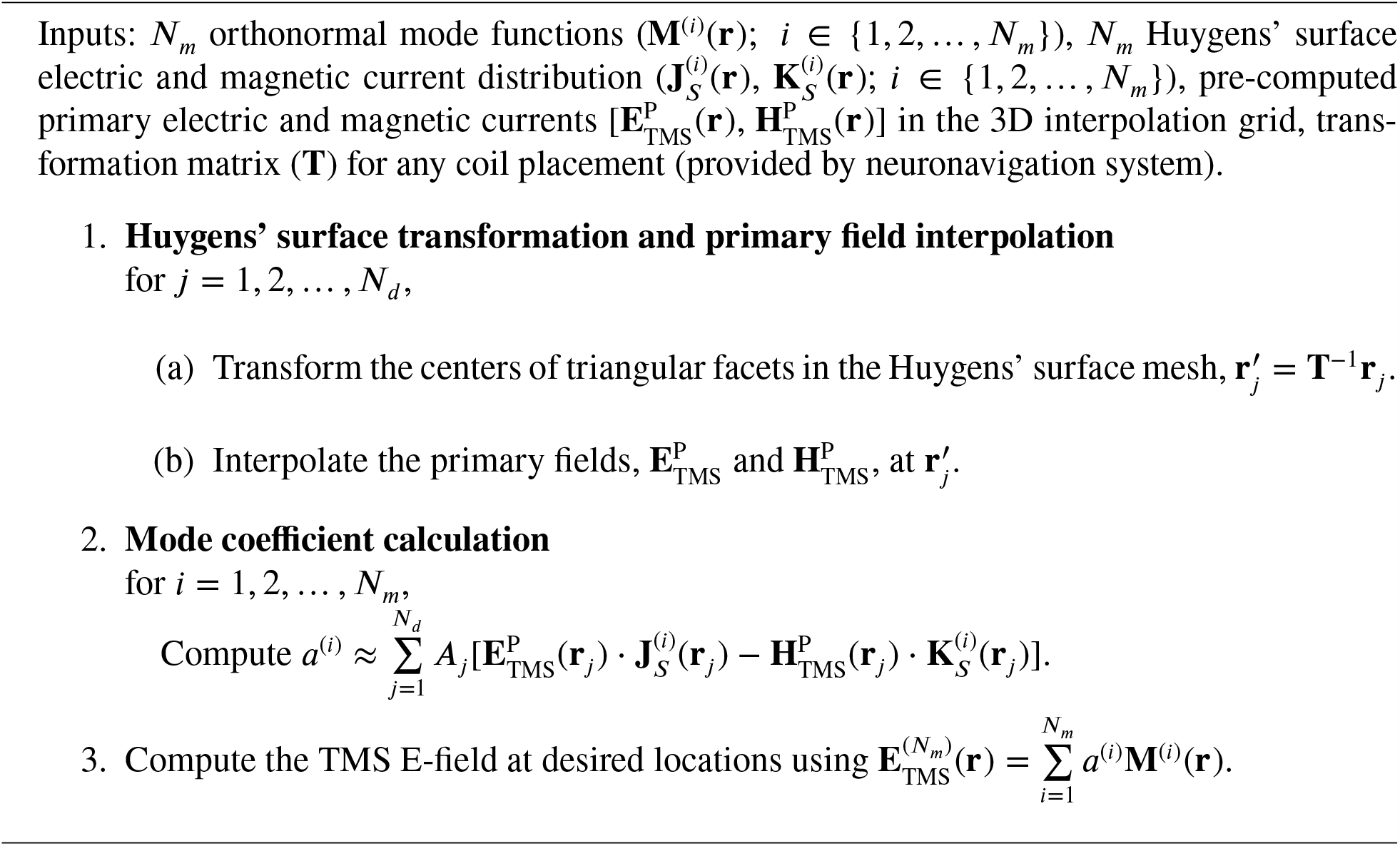

During the preprocessing stage (offline stage or, mode calculation stage), we compute the E-fields in the brain (GM and WM) consisting of 1.31–1.84 million tetrahedrons for ‘headreco’ models and 1.55–1.65 million tetrahedrons for ‘mri2mesh’ models. During the real-time stage, we compute the E-field at the barycenter of each triangular facet on the middle GM surface (a surface approximately midway into the GM, [19]) consisting of 122,000–289,000 triangular elements for ‘headreco’ models and 241,000–284,000 triangular elements for ‘mri2mesh’ models.

Our method is suitable for modeling any TMS coil; to illustrate, we included different coil types and sizes in this study. The Figure-8 coil consists of two 9 turn concentric circular loops with inner and outer loop diameters of 53 mm and 88 mm, respectively, which matches the 70-mm Figure-8 #31 in [39] and is approximated with the coil model described in [30]. The circular and double cone coil models were obtained from [40] and are models of the MagVenture Cool-40 Rat coil and D-B80 coil, respectively.

### 2.7. Error metrics

We performed a benchmark comparison between our real-time algorithm with the conventional 1^st^-order FEM solver. We consider global vector error (GVE) and global magnitude error (GME) as means of comparison between the FEM-calculated E-field (**E**_TMS_(**r**)) and the real-time computed E-field 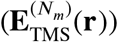, defined as follows:

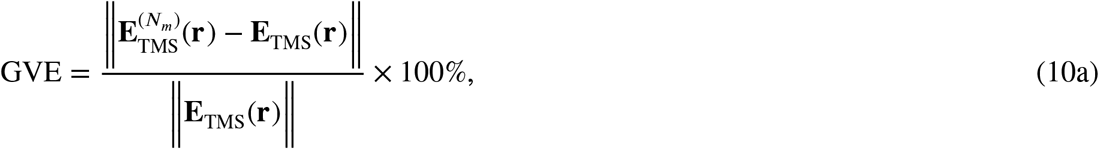

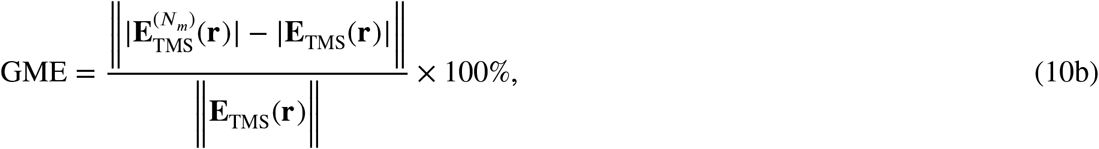

where ‖⋅‖ denotes the *L*^2^ norm defined in Section 2.1 and |⋅| is the magnitude for the E-field. Additionally, for a visual comparison of the E-fields, we consider the pointwise relative errors: local vector error (LVE) and local magnitude error (LME), normalized by the largest E-field magnitude in the ROI from the FEM solver defined as follows:

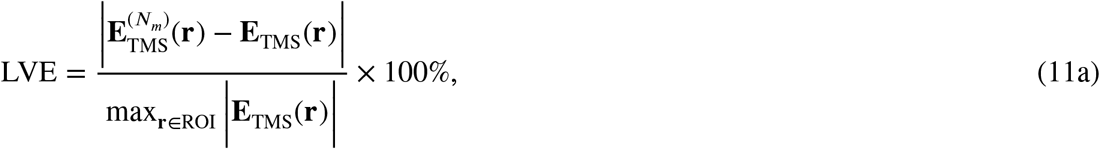

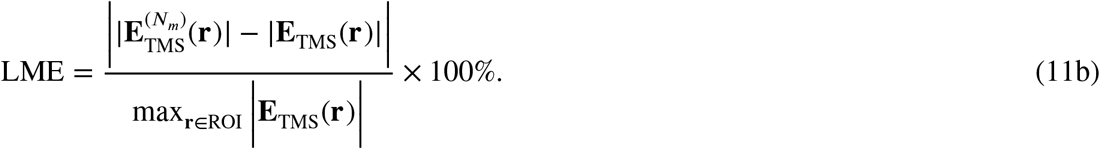

For all of our analyses, we perform simulations for modes 100 to 500, with a step size of 50.

## 3. Results

### 3.1. Accuracy of real-time predicted E-fields as a function of modes

In this section, we compare observed errors for all 16 head models and the 70-mm Figure-8 coil model. We randomly select 1000 coil placements for each head model and calculate the errors by comparing them with the reference 1^st^-order FEM solution. The convergence of GME and GVE is shown in Fig. 5 as a function of the number of modes. For ‘mri2mesh’ models, the mean GME and mean GVE are below 2% at modes (equals matrix rank) of 325 and 450, respectively. For ‘headreco’ models, the required modes are 350 and 475 for mean GME and GVE, respectively. There are some outlier errors, primarily corresponding to specific coil placements, which nonetheless remain below 3% for GVE and under 2% for GME at rank 500. Here we used the default ‘headreco’ and ‘mri2mesh’ mesh models whose FEM solution is known to have a GVE near 5% [37]. With 400 modes we observed a maximum GVE and GME error of 4% and 3%, respectively, across all simulations. To ensure that the real-time results are just as accurate as the 1^st^-order FEM, we estimated the error of the 1^st^-order FEM and real-time solutions by using a 2^nd^-order FEM as reference (results are given in the supplementary document). For 400 modes, the resulting difference in GVE and GME between real-time and 1^st^-order FEM was, on average, 0.17% and 0.14%, respectively [Fig. S3]. Furthermore, the GVE and GME across the 16,000 simulations showed that the real-time solution with 400 modes is up to 1.3% (1.7%) and 0.7% (1.1%) more (less) in agreement, respectively, than the 1^st^-order FEM to the 2^nd^-order FEM solution [Fig. S4 & S5], indicating that the real-time results are as accurate as the 1^st^-order FEM ones.

**Figure 5:**
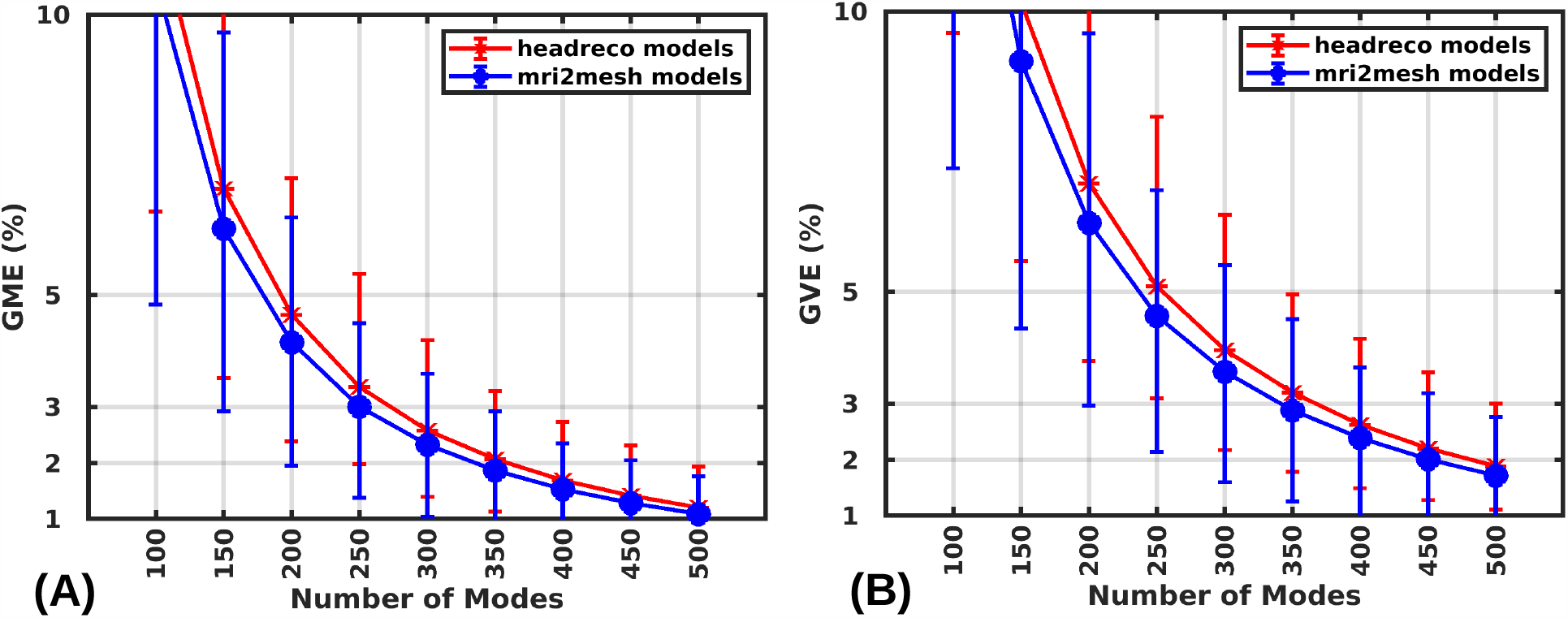
Convergence of global magnitude error (GME) (A) and global vector error (GVE) (B) as a function of the number of modes for both ‘mri2mesh’ and ‘headreco’ models with a 70-mm Figure-8 coil model. The error distribution for any mode is calculated across 8000 random coil placements (1000 random coil placements over the scalp of each of the 8 head models).

### 3.2. Effect of coil model on error convergence

Here consider the relative accuracy performance of the method for three distinct coil models 70-mm Figure-8, MagVenture D-B80 coil, and Cool-40 Rat coil. We consider the sample mean error over 1000 random coil placements over the scalp on each of the 16 head models and use the 1^st^-order FEM solution as a reference. The mean GME and the mean GVE are shown in Fig. 6. The mean GME is below 2% for ‘mri2mesh’ (as well as ‘headreco’) head models at ranks above 325, 425, and 375 (350, 375, and 375) for the Figure-8, D-B80, and Cool-40 coils, respectively. Due to the unique bending shape, the D-B80 coil has an E-field that has more fine features relative to the others, thereby, requiring comparatively more modes for its expansion. The mean GVE is below 2% for ‘mri2mesh’ (‘headreco’) head models at ranks above 450, 550, and 475 (475, 525, and 500) for the Figure-8, D-B80, and Cool-40 coils, respectively. All coils exhibit similar errors. However, compared to the D-B80 and Cool-40 coils, the Figure-8 coil model converges to the 2% error limit with fewer modes.

**Figure 6:**
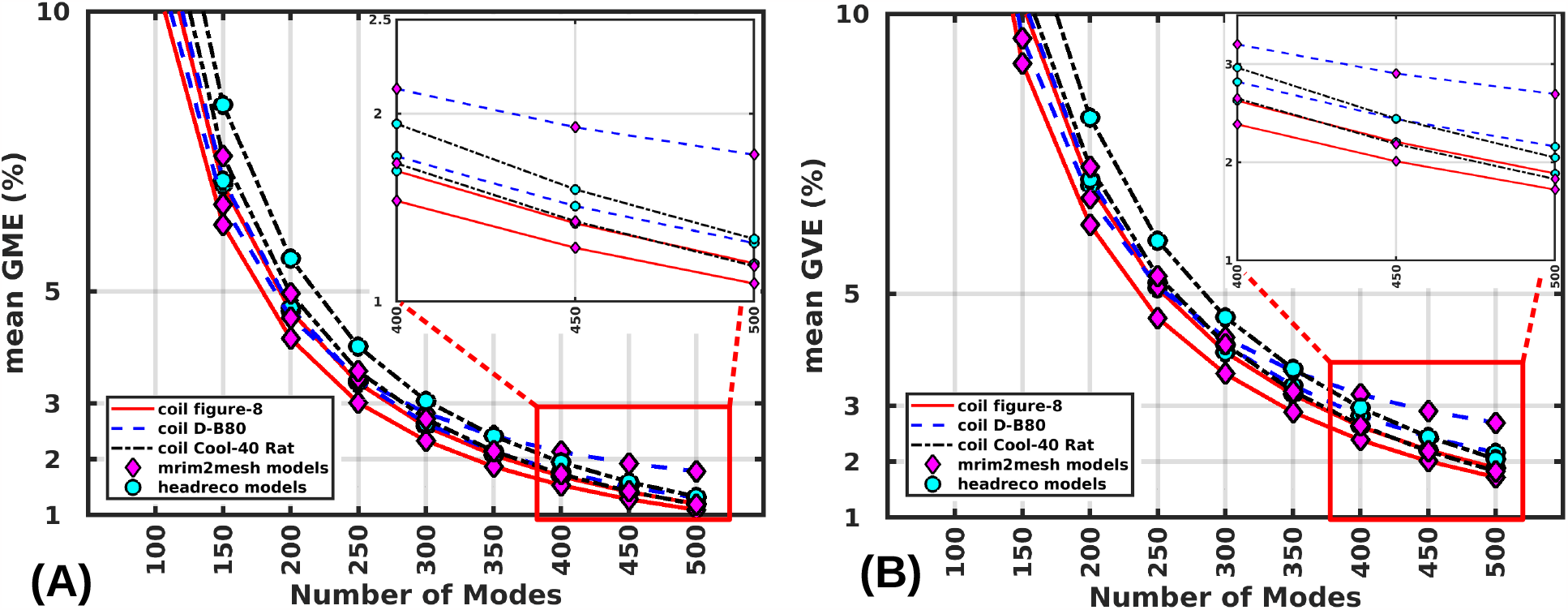
Convergence of mean global magnitude error (GME) in (A) and global vector error (GVE) in (B) as a function of the number of modes for both ‘mri2mesh’ and ‘headreco’ models with 3 coils (70-mm Figure-8, MagVenture D-B80 coil, and Cool-40 Rat coil). The mean error for any mode is calculated across 16,000 random coil placements (1000 random coil placements over the scalp of each of the 16 head models from 8 subjects). The inset of each plot shows the errors for the higher number of modes (400–500).

### 3.3. E-field visualization

In this section, several exemplary simulation results of coil placements over the scalp of the ‘Ernie’ head model are shown. Fig. 7 shows the specific coil placement over the scalp, the associated E-field induced on the middle GM surface computed in Real-time and FEM solvers, and the corresponding local magnitude error (LME) and local vector error (LVE) distributions. The E-field distributions predicted in real-time and FEM are visually indistinguishable. Furthermore, the peak E-field is the same up to 0.65 V⁄m in all cases shown. The maximum LME for each scenario (top to bottom) are 3.7%, 3.6%, 2.7%, 3.1%, 2.9%, and 3.2%, whereas the corresponding maximum LVE is 4%, 3.8%, 3.9%, 3.2%, 3.4%, and 4.5%, respectively. These results indicate that the real-time predicted E-field distributions are equally valid to the FEM 1^st^-order solutions, which have an LME of about 5%.

**Figure 7:**
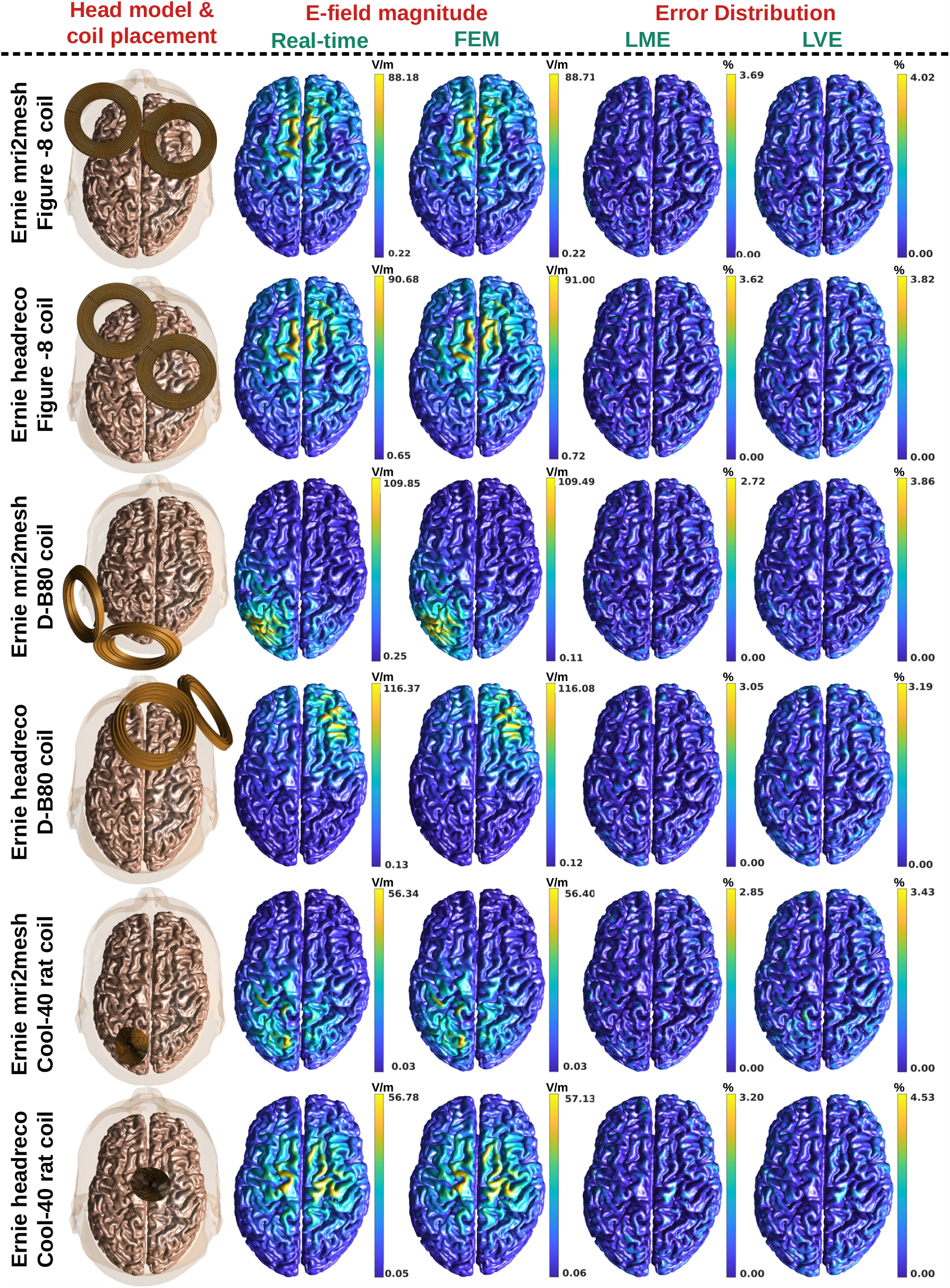
Illustration of the real-time TMS-induced E-field (2^nd^ column) and FEM-induced E-field (3^rd^ column) on the middle GM surface for randomly chosen coil placements (1^st^ column) over the scalp of SimNIBS 3.2’s ‘Ernie’ head model. The last two columns show the local error distributions (LME and LVE) over the middle GM surface.

### 3.4. Computational run-time and memory requirements

Here we consider single-precision arithmetic during real-time computation, as the results remained unchanged up to seven digits relative to double-precision. Fig. 8(A) and (B) show the mean mode calculation time (pre-processing time) as a function of the number of modes across 8 ‘mri2mesh’ models and 8 ‘headreco’ models, respectively. The pre-processing stage is computed using an AMD Rome 2.0 GHz CPU. On average the pre-processing time required to generate 400 modes is 38 hours and 34 hours for ‘mri2mesh’ and ‘headreco’ models, respectively. The total computational runtime is 2 FEM runs per mode. Our single-threaded implementation requires, on average, 3 minutes of computation time per simulation in an AMD Rome 2.0 GHz processor. Therequired pre-processing computations could be significantly sped up by using multiple threaded FEM solvers.

**Figure 8:**
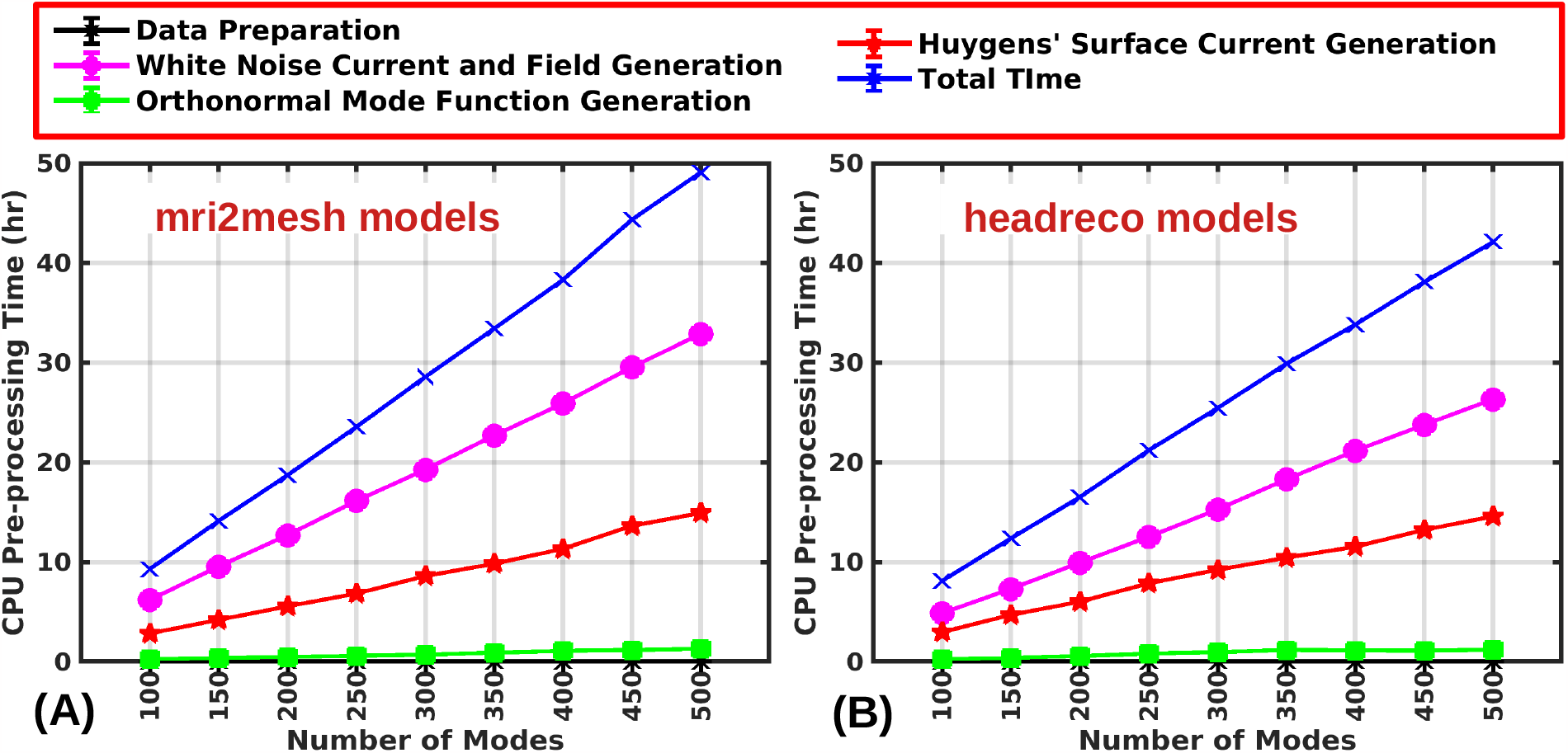
Mean computational time for pre-processing stage (mode and field generation stage) for ‘mri2mesh’ models (A) and ‘headreco’ models (B). At any rank (mode), the time is calculated across 8 head models from 8 subjects.)

Fig. 9 shows the computational reconstruction time as a function of the number of modes for a GPU and CPU, respectively. The above result was obtained by running 48000 simulations across the 16 head models and 3 coil models. The mean reconstruction time to predict the E-field over the intermediate GM-WM surface for a fixed coil placement is 2.2 ms for 400 modes in an NVIDIA RTX 3080 GPU with a maximum time of 3.8 ms . The first step of the real-time TMS (coordinate transformation of the Huygens’ surface) takes only 0.03 ms, whereas the CPU takes 0.9 ms (a 30 times speed-up). The second step (multi-linear interpolation of primary fields) takes 1.7 ms in a GPU and 37.40 ms in a CPU. In other words, the GPU requires 22 times less run-time than the CPU. The mode coefficient calculation is completed in 0.4 ms in a GPU and 1100 ms in a CPU (i.e., a 2750 times faster). Finally, the fourth step (TMS E-field computation) is rapidly computed in 0.03 ms in a GPU vs. 105 ms in the CPU (3500 times faster). Overall, the mean total time for estimating the TMS-induced E-field using 400 modes is 2.2 ms in a GPU and 1200 ms in the CPU (550 times faster). This ratio could be improved by accelerating the Huygens’ surface coordinate transformation.

**Figure 9:**
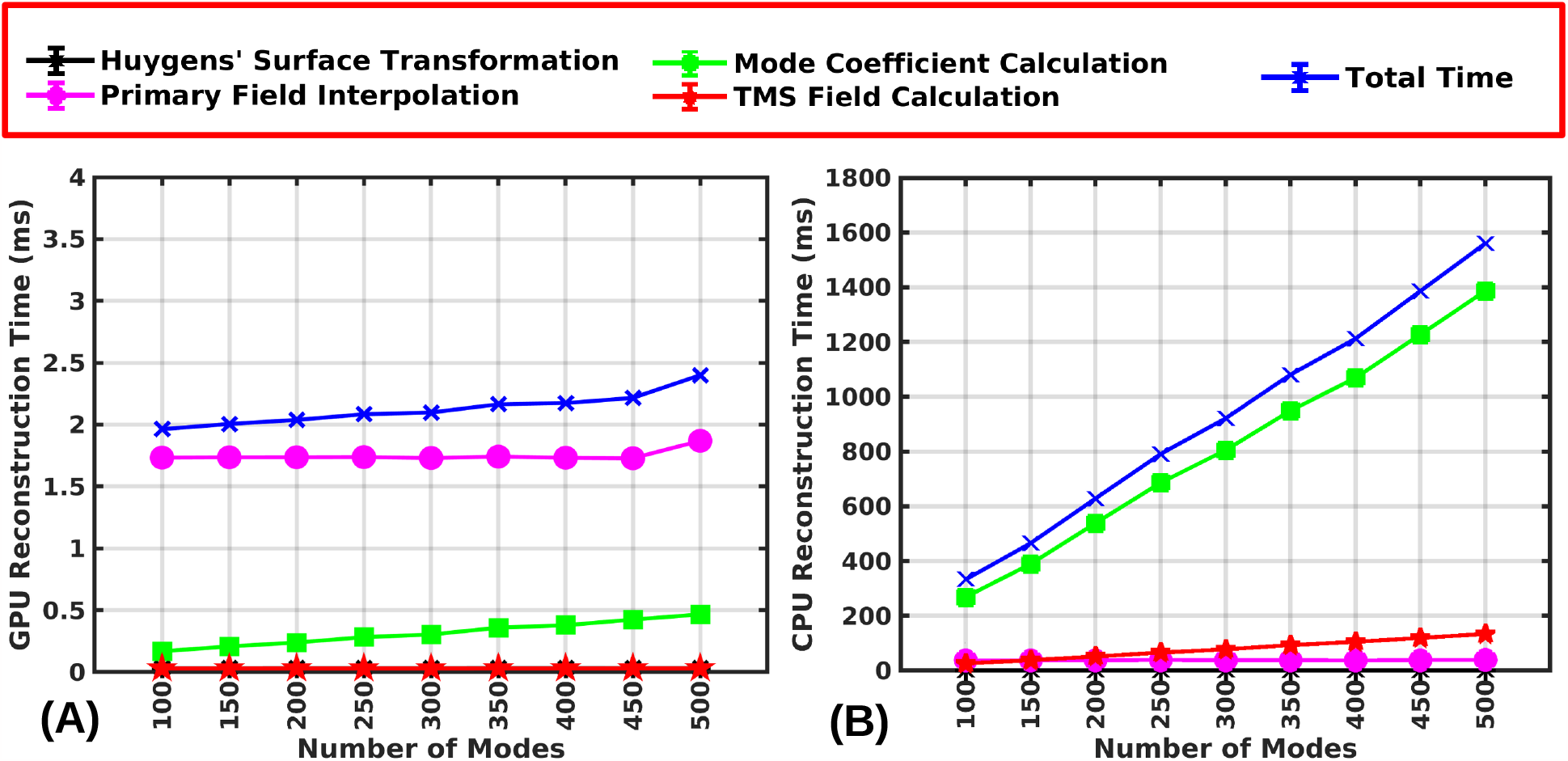
E-field reconstruction time in GPU (A) and CPU (B) as a function of the number of modes. For any mode, the time is calculated across 48,000 random coil placements (1,000 random coil placements over the scalp of each head model from each subject for each coil model.)

Fig. 10 shows the required memory in the reconstruction stage for both CPU (AMD Rome CPU, 2.0 GHz) and GPU (NVIDIA RTX 3080-10 GB) across 14 head models. The total required memory in the reconstruction stage is the same for both the GPU and the CPU. The differences in GPU and CPU memory requirements stem from the fact that the Matlab environment requires overhead that is not accounted for in the GPU memory. In other words, the GPU only has all required data structures (e.g., modes, surface currents, and interpolatory primary fields). When the real-time computation is performed in the GPU, the required mean CPU and the GPU memory for 400 modes are 1.3 GigaBytes(GB) and 3 GB, respectively. Additionally, the required mean CPU memory during real-time computation in the same CPU is 4.3 GB .

**Figure 10:**
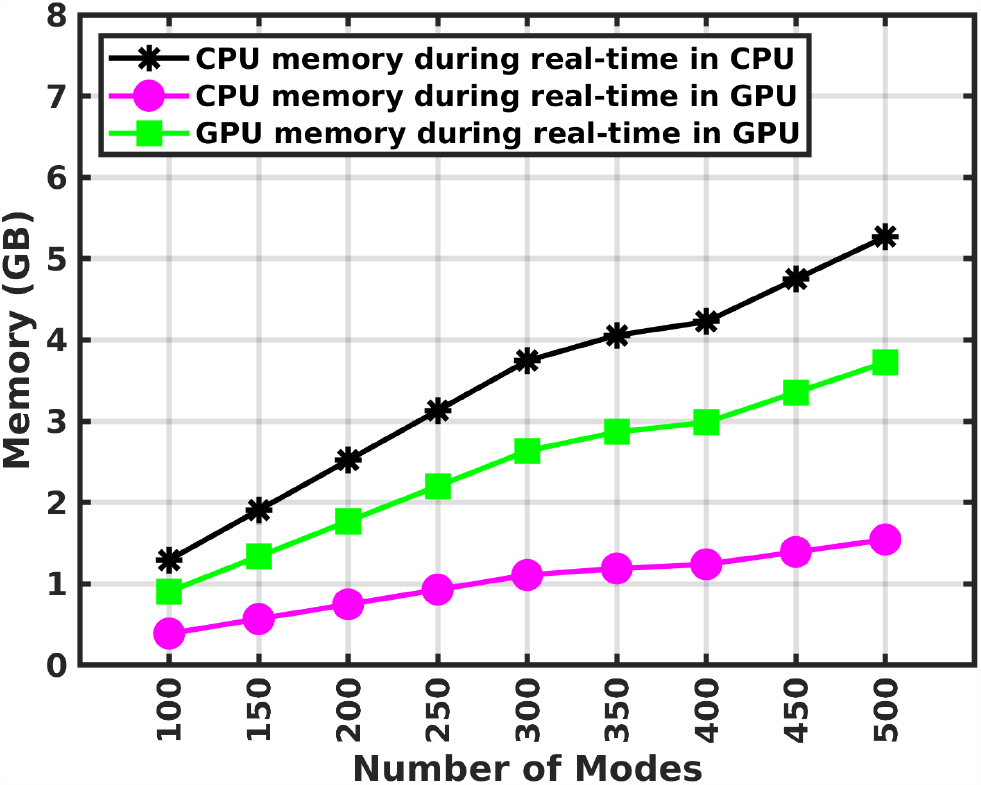
Mean computational memory (in Gigabytes, GB) requirement in a CPU and GPU as a function of the number of modes when the real-time operations are done in a CPU or GPU.

## 4. Discussion

The real-time TMS E-field approximation method excels in rapidly calculating the E-field over the middle GM surface using GPU acceleration, achieving this computation within 4 ms for any coil placement over the scalp of the subject. Our detailed benchmarking analysis indicates that 450 modes are enough to achieve a mean GME and a mean GVE below 2%. Additionally, with only 400 modes, the maximum GME and maximum GVE are 2.8% and 4.2%, respectively. Therefore, with 400 modes, the agreement between the real-time predicted E-fields and 1^st^-order FEM is closer than the expected 5% error of the FEM solution. These results are consistent across multiple subjects, head model construction pipelines, coil types, and coil placements.

In this study, we conducted a benchmark comparison between our real-time algorithm and the 1^st^-order FEM solver, which has a known relative error of about 5% [34]. Correspondingly, we observed that the convergence of the mode expansion greatly decelerated below 5%. This is likely because, below the FEM solution error threshold, we are increasingly spanning the erroneous part of the solution, which is not expected to be spanned by a small number of modes. We additionally used 2^nd^-order FEM, which results in a more accurate E-field prediction, for a single subject to generate the mode expansion. The results given in the supplement (Fig. S3, S4, and S5) indicate that using a more accurate solver could result in slightly improved convergence. At the current time, running the 2^nd^-order FEM requires excessive preprocessing time, thereby limiting the applicability of this method. Although not pursued here, this tool could be implemented using efficient implementations of BEM [41] that are more accurate than 1^st^-order FEM [30] to achieve improved convergence. Furthermore, 1^st^-order FEM is currently the most commonly used tool for TMS E-field dosimetry.

Our method relies on the existence of a small set of modes that span the possible range of E-fields induced by a TMS coil. From spherical solutions, we expect that the total required modes are larger for superficial regions relative to deeper ones. As such, this method could in principle be applied to determine E-fields on more superficial compartments like the CSF, skull, and skin. However, a larger number of modes would likely be necessary in such cases, which could potentially limit its applicability.

The preprocessing stage can be done using any available FEM or BEM code. Using our 1^st^-order in-house FEM [30, 32], which is optimized for accuracy, requires less than 40 hours for 400 modes at 180 s per FEM run. However, a faster implementation of the underlying FEM can speed up the process. For example, the SimNIBS implementation of FEM requires a runtime of 29.8 s, which reduces the total time to 6.6 hours.

The real-time stage requires only under 4 ms (assuming 400 modes) in an NVIDIA RTX 3080 GPU and 1.2 s in an AMD Rome 2.0 GHz CPU to predict the TMS-induced E-field. The reconstruction time is almost independent of the number of modes in a GPU. Therefore, for better accuracy, a higher number of modes can be accommodated in a higher-end GPU.

We found empirically that the pre-processing stage should be computed in double-precision arithmetic whereas the real-time stage can be performed in single-precision arithmetic for accommodating more modes inside the GPU and a larger ROI (i.e., the whole brain/head). A GPU with only 4 GB of dedicated memory is large enough for this process accommodating up to 400 modes, pre-computed primary fields, Huygens’ surface nodes, and an ROI of size 220,000 samples on the middle GM surface.

Future work includes connecting this GPU-based E-field computation and GPU-based visualization within one screen refresh in real-time. The integration of real-time E-field modeling in combination with neuronavigation and TMS stimulator control will enable several avenues of research and clinical intervention. For example, this methodology will enable support for more reliable dosing throughout a TMS session by automatically changing TMS intensities accounting for coil placement variability. It will also allow the development of methods for faster motor threshold determination applying fewer TMS pulses by using E-field-based instead of grid search approaches. Furthermore, this approach will allow the development of rapid coil placement optimization for updated brain targets and E-field constraints during the TMS procedure. Updated brain targeting may be derived from behavioral (e.g., task performance, or treatment indicators) or physiological (e.g., heart rate variability, or electroencephalography) response even in closed-loop settings.

## 5. Conclusions

We have developed a method for rapidly calculating the TMS-induced E-field. This method is based on a functional generalization of the probabilistic matrix decomposition (PMD) and generalized Huygens’ and reciprocity principles. The initial preprocessing stage takes approximately 40 hours to complete. However, subsequent E-field computations can be done within 4 ms in a GPU and 1.2 s in a CPU over the middle GM surface (containing, on average, 216,000 barycentric points). The resultant E-field has accuracy comparable to standard FEM solutions. Notably, this computational performance can be achieved using a standard GPU with a dedicated memory of only 4 GB, making it practical for many users. This framework enables real-time E-field calculation in the cortex for arbitrary coil design and coil placement, and generalizes well for distinct head model pipelines, underscoring its adaptability and suitability for a wide range of applications.

**Table 1.** Abbreviation and notation.

| Notation | Definition |
| --- | --- |
| $N_m$ | number of modes |
| $N_e$ | number of brain tetrahedrons |
| $N_d$ | number of triangular facets in the Huygens' surface mesh |
| $I'(t)$ | time derivative of the driving current pulse waveform during TMS |
| $\mathbf{W}^{(i)}(\mathbf{r}_j)$ | $i^{\text{th}}$ sample of white noise current at centers of $j^{\text{th}}$ triangular facet ($j \in \{1, 2, \dots, N_d\}$) in the Huygens' surface; $i \in \{1, 2, \dots, N_m\}$ |
| $\mathbf{E}_W^{P(i)}(\mathbf{r})$ | primary E-field induced in the brain due to current source $\mathbf{W}^{(i)}(\mathbf{r}_j)$; $i \in \{1, 2, \dots, N_m\}$ |
| $\mathbf{E}_W^{(i)}(\mathbf{r})$ | total induced E-field in the brain due to current source $\mathbf{W}^{(i)}(\mathbf{r}_j)$; ($i \in \{1, 2, \dots, N_m\}$) |
| $\phi^{(i)}(\mathbf{r})$ | scalar potential in the brain; $i \in \{1, 2, \dots, N_m\}$ |
| $\mathbf{M}^{(i)}(\mathbf{r})$ | $i^{\text{th}}$ orthonormal mode function |
| $\mathbf{E}_M^{(i)}(\mathbf{r}), \mathbf{H}_M^{(i)}(\mathbf{r})$ | E-field and H-field generated due to $i^{\text{th}}$ impressed current (mode function) $\mathbf{M}^{(i)}(\mathbf{r})$; $i \in \{1, 2, \dots, N_m\}$ |
| $\mathbf{J}_S^{(i)}(\mathbf{r}), \mathbf{K}_S^{(i)}(\mathbf{r})$ | electric and magnetic current on the Huygens' surface, $S$, for $i^{\text{th}}$ mode function; $i \in \{1, 2, \dots, N_m\}$ |
| $\mathbf{E}_{J_S, K_S}^{(i)}(\mathbf{r})$ | E-field generated due to $\mathbf{J}_S^{(i)}(\mathbf{r})$ and $\mathbf{K}_S^{(i)}(\mathbf{r})$; $i \in \{1, 2, \dots, N_m\}$ |
| $\mathbf{E}_{\text{TMS}}^P(\mathbf{r}), \mathbf{H}_{\text{TMS}}^P(\mathbf{r})$ | primary TMS E-field and H-field due to any coil model |
| $\mathbf{T}$ | transformation matrix for relative placement of Huygens' surface with respect to the coil |
| $a^{(i)}$ | mode coefficient corresponding to $i^{\text{th}}$ mode function ($\mathbf{M}^{(i)}(\mathbf{r})$); $i \in \{1, 2, \dots, N_m\}$ |
| $\mathbf{J}_{\text{TMS}}(\mathbf{r})$ | magnetic current on the TMS coil |
| $\mathbf{E}_{\text{TMS}}(\mathbf{r})$ | actual TMS induced E-field |
| $\mathbf{E}_{\text{TMS}}^{(N_m)}(\mathbf{r})$ | real-time approximated TMS induced E-field from $N_m$ mode functions |
| $A_j$ | area of the $j^{\text{th}}$ triangular facet in the Huygens' surface; $j \in \{1, 2, \dots, N_d\}$ |
| $V_k$ | volume of the $k^{\text{th}}$ tetrahedron in the head mesh; $k \in \{1, 2, \dots, N_e\}$ |
| $\mathbf{R}_c$ | $3 \times 3$ rotation matrix for TMS coil with respect to the head |
| $\mathbf{T}_0$ | translation vector for the TMS coil placement with respect to the head |
| $\sigma(\mathbf{r})$ | conductivity at location $\mathbf{r}$ in the head |
| $\hat{\mathbf{n}}$ | normal vector on the scalp surface pointing outward |
| $S$ | Huygens' surface |
| $\Omega$ | brain region in the head |

## Appendix

### CRediT authorship contribution statement

**Nahian I. Hasan:** Conceptualization, Methodology, Software, Validation, Formal analysis, Investigation, Data curation, Writing - original draft, Writing - review & editing, Visualization. **Moritz Dannhauer:** Validation, Writing - review & editing. **Dezhi Wang:** Writing - review & editing. **Zhi-De Deng:** Supervision, Writing - review & editing. **Luis J. Gomez:** Funding acquisition, Conceptualization, Supervision, Methodology, Software, Validation, Formal analysis, Investigation, Writing - review & editing, Funding acquisition.

### Declaration of competing interest

None

### Funding

Research reported in this publication was supported by the National Institute of Mental Health of the National Institutes of Health under Award Number R00MH120046. M. Dannhauer and Z.- D. Deng are supported by the National Institute of Mental Health Intramural Research Program (ZIAMH002955). The content of the current research is solely the responsibility of the authors and does not necessarily represent the official views of the National Institutes of Health.

### Data and Code Availability Statement

Data and Code Availability Statement: In-house codes are openly available at this link - **Link will be available after publication**

### Supplementary Material

Supplementary materials can be found in the online version of this research article.

