## supplementary PDF for "Real-Time Computation of Brain E-Field for Enhanced Transcranial Magnetic Stimulation Neuronavigation and Optimization"

### Supplementary Material

#### S1.1. Effect of interpolation grid density

Fig. S1 shows the relative error (GME) with respect to the interpolation grid density for 1000 random coil placements over the scalp of a subject when the number of modes is fixed at 400. There is negligible change in the performance until a grid spacing of 5 mm in the order of $10^{-6}$. On the other hand, at extremely low grid dimensions, the required memory increases which leaves less room for the modes to be accommodated in the GPU memory.

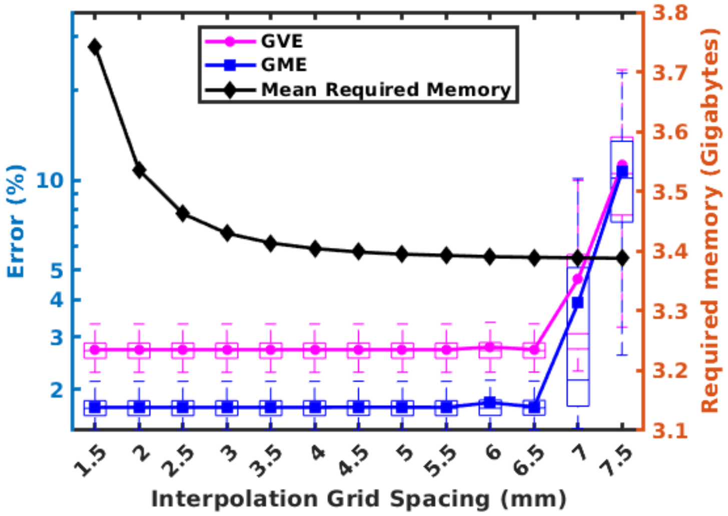


**Figure S1:** Effect of interpolation grid density on real-time TMS E-field prediction. The average error remains almost constant until a grid spacing of 5 mm. On the right axis, the effect on memory is shown. For lower grid dimensions (denser grid), the required memory increases rapidly.

#### S1.2. Effect of FEM order

Fig. S2 shows the effect of FEM order on the convergence of error. For higher FEM orders, the convergence is slower in both GME and GVE. For example, we conducted this study on the 'Ernie' head model from SimNIBS. For first-order FEM, the GME and GVE reach the 2% error bound at 360 and 480, respectively. On the other hand, for second-order FEM, the same error bound is reached at 290 and 330, respectively. But, the trade-off is the time and memory requirement in the offline stage. The real-time stage is completely unaffected by the order of the FEM. However, if the targeted error limit is very high, the improvement becomes negligible.

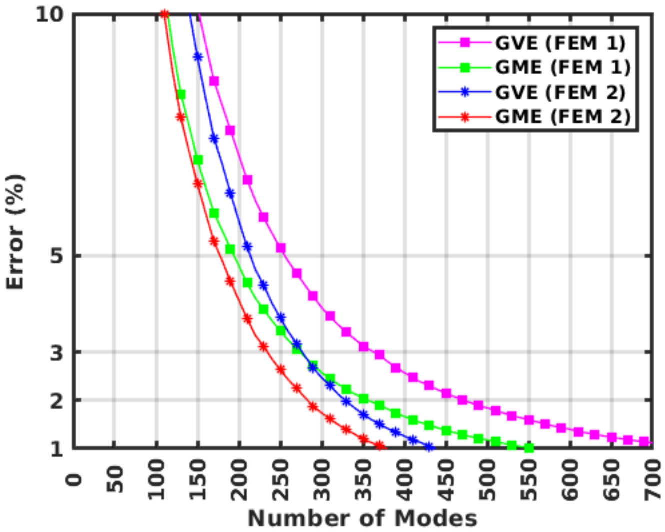


**Figure S2:** Effect of FEM on the error of real-time TMS E-field prediction. Higher order FEM facilitates less number of modes for a targeted error bound.

#### S1.3. Accuracy of E-field due to real-time TMS and 1$^{st}$-order FEM

Here, we show the relative accuracy of the real-time predicted TMS E-field and the 1$^{st}$-order FEM with respect to the 2$^{nd}$-order FEM. Fig. S3 shows the difference between the $GVE_{RT}$ (GVE between the real-time TMS and 2$^{nd}$-order FEM) and the $GVE_{FEM}$ (GVE between the 1$^{st}$-order FEM and 2$^{nd}$-order FEM) along with the corresponding differences in GMEs. At rank 400, the real-time predicted E-field almost perfectly matched the 1$^{st}$-order FEM accuracy, beyond which the algorithm is likely to model the noise. Therefore, the real-time algorithm is as accurate as the underlying 1$^{st}$-order FEM solver. Fig. S4 and Fig. S5 show the distributions of errors (GVE & GME, respectively) differences between the real-time E-field and the 1$^{st}$-order FEM with respect to the 2$^{nd}$-order FEM.

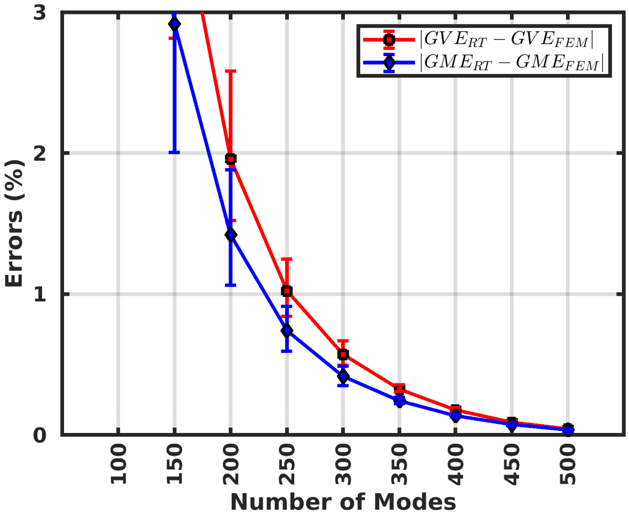


**Figure S3:** Error (GVE & GME) comparison between real-time TMS E-field and $1^{st}$-order FEM-induced E-field with respect to the $2^{nd}$-order FEM-induced E-field. The distribution of error differences at any rank (mode) is calculated from 16000 random simulations (1000 random coil placements over the scalp of each of 16 head models).

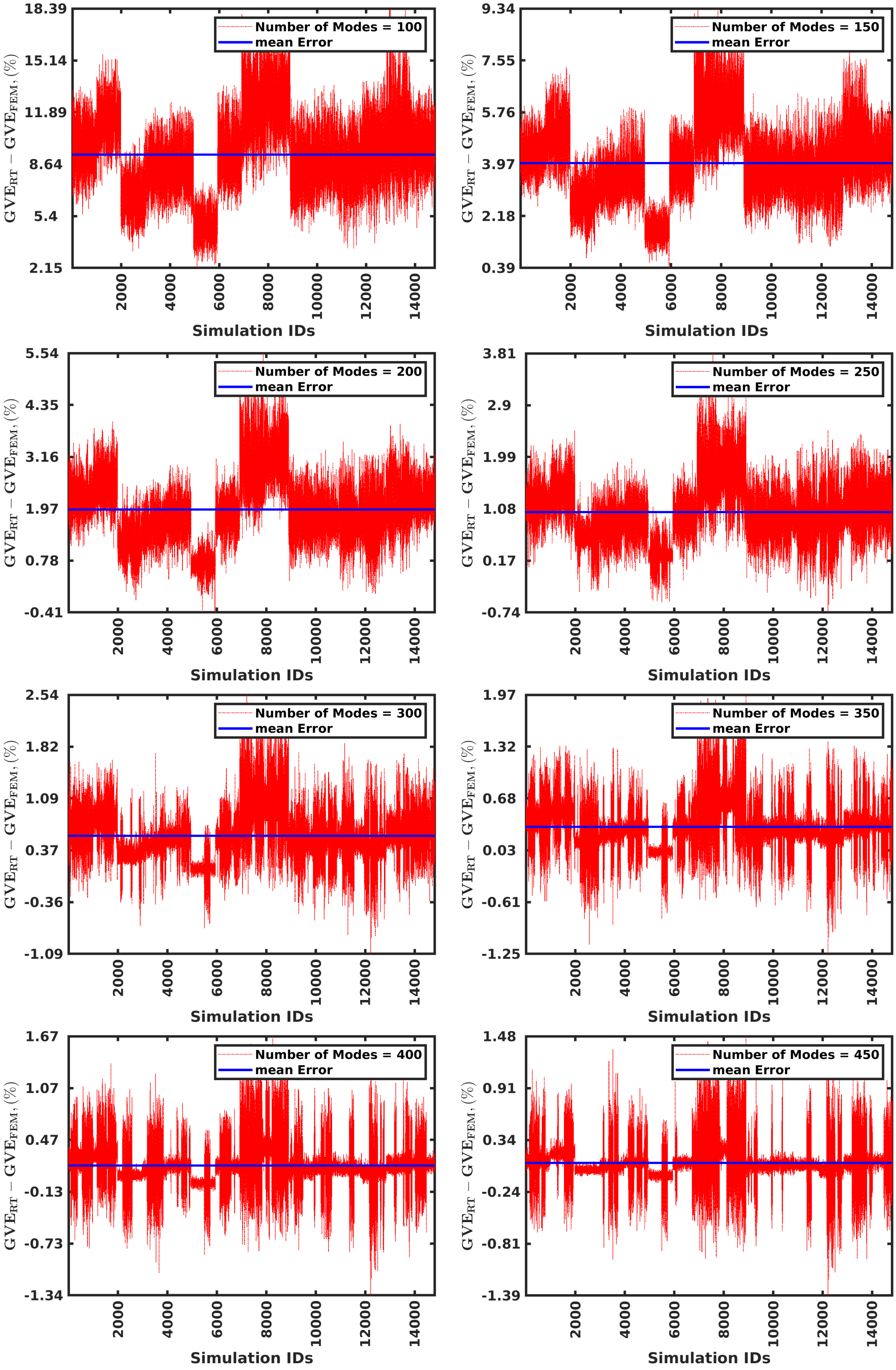


**Figure S4:** Error (GVE) comparison between real-time TMS E-field and $1^{st}$-order FEM-induced E-field with respect to the $2^{nd}$-order FEM-induced E-field across 16000 simulations for the ranks of 100, 150, 200, 250, 300, 350, 400, and 450.

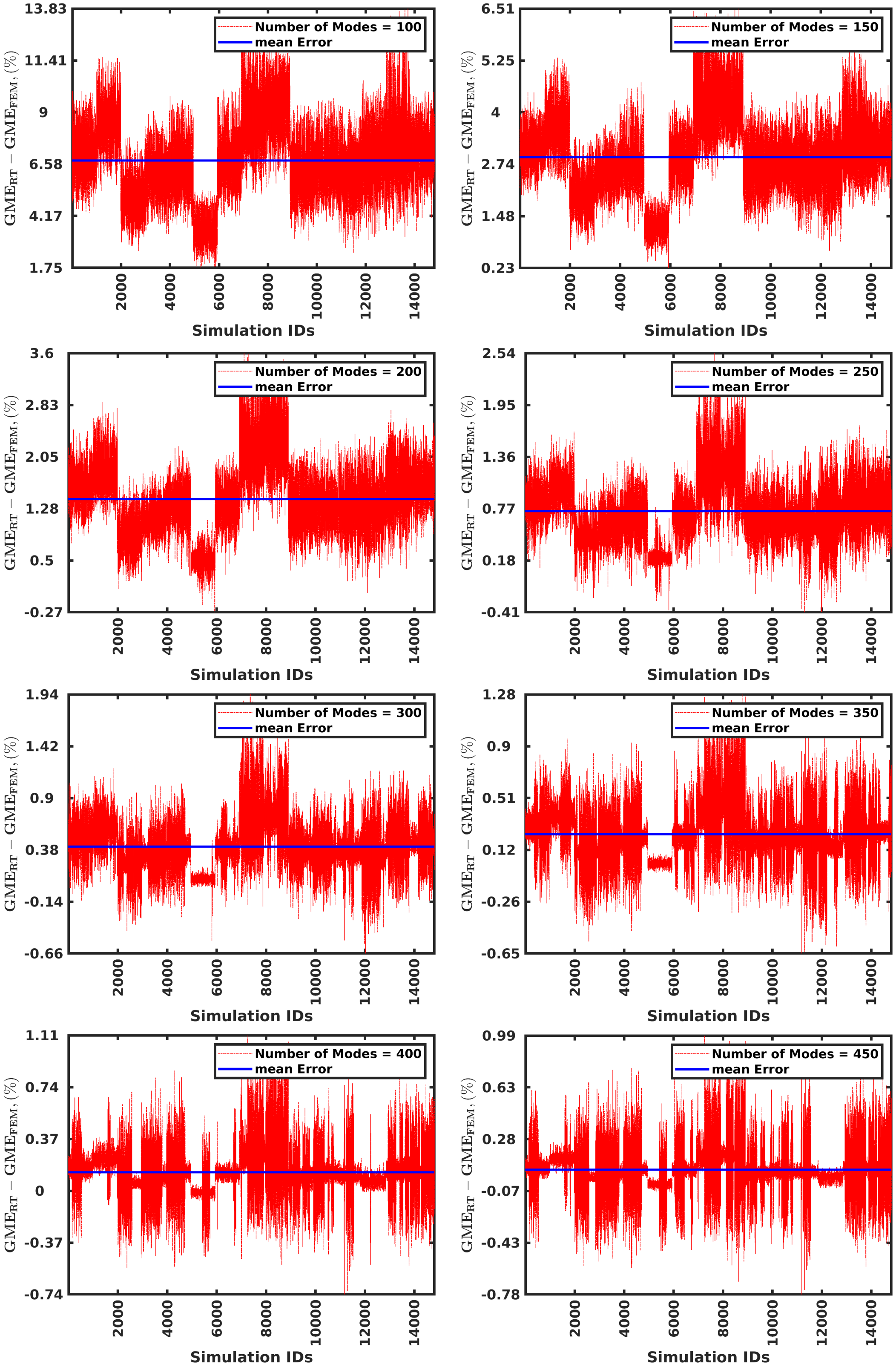


**Figure S5:** Error (GME) comparison between real-time TMS E-field and $1^{st}$-order FEM-induced E-field with respect to the $2^{nd}$-order FEM-induced E-field across 16000 simulations for the ranks of 100, 150, 200, 250, 300, 350, 400, and 450.
